# Spatial and temporal localization of cell wall associated pili in *Enterococcus faecalis*

**DOI:** 10.1101/2021.07.23.451969

**Authors:** Pei Yi Choo, Charles Y. Wang, Michael S. VanNieuwenhze, Kimberly A. Kline

## Abstract

*Enterococcus faecalis* relies upon a number of cell wall-associated proteins for virulence. One virulence factor is the sortase-assembled endocarditis and biofilm associated pilus (Ebp), an important factor for biofilm formation *in vitro* and *in vivo*. The current paradigm for sortase-assembled pilus biogenesis in Gram-positive bacteria is that the pilus sortase covalently links pilus monomers prior to recognition, while the housekeeping sortase cleaves at the LPXTG motif within the terminal pilin subunit, and subsequently attaches assembled pilus fiber to the growing cell wall at sites of new cell wall synthesis. While the cell wall anchoring mechanism and polymerization of Ebp is well characterized, less is known about the spatial and temporal deposition of this protein on the cell surface. We followed the distribution of Ebp and peptidoglycan (PG) throughout the *E. faecalis* cell cycle via immunofluorescence microscopy and fluorescent D-amino acids (FDAA) staining. Surprisingly, cell surface Ebp did not co-localize with newly synthesized PG. Instead, surface-anchored Ebp was localized to the cell hemisphere but never at the septum where new cell wall is deposited. In addition, the older hemisphere of the *E. faecalis* diplococcus were completely saturated with Ebp, while Ebp appeared as two foci directly adjacent to the nascent septum in the newer hemisphere. A similar localization pattern was observed for another cell wall anchored substrate by sortase A, aggregation substance (AS), suggesting that this may be a general rule for all SrtA substrates in *E. faecalis*. When cell wall synthesis was inhibited by ramoplanin, an antibiotic that binds and sequesters lipid II cell wall precursors, new Ebp deposition at the cell surface was not disrupted. These data suggest an alternative paradigm for sortase substrate deposition in *E. faecalis*, in which Ebp are anchored directly onto un-crosslinked cell wall, independent of new PG synthesis.

## Introduction

*Enterococcus faecalis* is a Gram-positive ovococcal bacteria commonly found in the gastrointestinal tracts of humans and other mammals. *E. faecalis* is a public health concern due to its resistance to many antimicrobial drugs (Arias and Murray, 2012). *E. faecalis* is an opportunistic pathogen capable of causing life-threatening infections in humans such as infective endocarditis, bacteremia, and urinary tract infection (UTI) (Hidron et al., 2008; Murdoch et al., 2009; Patterson et al., 1995; Weiner-Lastinger et al., 2019). Its ability to colonize and infect the human host involves multiple virulence factors which are responsible for adhesion and biofilm formation. These virulence factors include aggregation substance (AS), collagen binding protein (Ace), enterococcal surface protein (Esp), and endocarditis and biofilm-associated pilus (Ebp) (Hendrickx et al., 2009). Unlike Gram-negative bacteria that possess an outer membrane, the Gram-positive bacterium *E. faecalis* contains a thick peptidoglycan (PG) cell wall, which acts as a scaffold for the processing and attachment of these virulence factors. A feature common to each of these cell-wall associated virulence factors is the C-terminal cell wall sorting (CWS) signal which consists of an LPXTG motif, a hydrophobic domain, and a positively charged cytoplasmic tail (Schneewind et al., 1992).The enzyme responsible for the recognition and anchoring of these surface proteins to the peptidoglycan is the housekeeping enzyme, sortase A (SrtA) (Mazmanian et al., 1999; Ton-That et al., 1999). SrtA-mediated sorting of virulence factors to the cell wall is crucial to *E. faecalis* pathogenesis and long term persistence.

Ebp are well-characterized cell wall-attached surface protein in *E. faecalis*. Ebp are important for biofilm formation and are implicated in endocarditis and UTI (Nallapareddy et al., 2011; Sillanpaa et al., 2013). Each Ebp is composed of three pilin subunits (EbpA, EbpB, and EbpC), where EbpA and EbpB are minor subunits that form the pilus tip and base, respectively, while EbpC is the major subunit that makes up the pilus backbone (Nallapareddy et al., 2006; Nielsen et al., 2013). Ebp encoding genes are in an operon together with sortase C (*srtC*), which is the sortase responsible for Ebp subunit polymerization (Nallapareddy et al., 2011). Assembled pili are then anchored onto the cell wall by SrtA. Through this sorting mechanism, fully polymerized and cell wall-anchored Ebp become surface-exposed and can facilitate adhesion to abiotic and biotic surfaces.

SrtA is a membrane-anchored transpeptidase conserved in most Gram-positive bacteria (Marraffini et al., 2006). SrtA is focally enriched at the septum of *E. faecalis* cells (Kline et al., 2009). SrtA substrates are translocated across the cell membrane by the general Sec secretion machinery, whereupon they are transiently membrane-associated with the cell membrane via their C-terminal transmembrane domain. An LPXTG motif within the membrane-associated substrates is recognized and cleaved by the sortase enzyme between the threonine and glycine residue. Cleavage results in a covalent linkage between the active site cysteine residue of SrtA and the carbonyl group of the threonine residue of the substrate to generate an acyl-enzyme intermediate. This intermediate is then covalently captured by a lipid II cell wall precursor and is subsequently incorporated into the growing peptidoglycan layer via the well-established transglycosylation and transpeptidation reactions (Perry et al., 2002; Ruzin et al., 2002; Ton-That and Schneewind, 1999). The mechanism of SrtA has largely been defined in *Staphylococcus aureus*, which has served as a model organism for Gram-positive SrtA function. Although there is strong evidence that the crossbridge on lipid II is the anchoring site for sortase substrates, and the prevailing model is that the lipid II precursor is the SrtA target, the possibility of substrate anchorage onto the crossbridge of older uncrosslinked PG has not been eliminated.

The Enterococcal cell wall, like all Gram-positive bacteria, is composed mainly of PG, wall teichoic acid, and lipoteichoic acid (Rajagopal and Walker, 2017). The PG layers sit directly above the phospholipid bilayer, forming a lattice structure that protects the cell from osmotic stress and pressure. A single *E. faecalis* PG unit is made up of disaccharide N-acetylmuramic acid-N-acetylglucosamine (NAM-NAG) with a pentapeptide stem attached to NAM and an L-Ala-L-Ala crossbridge attached to the ε-amino group of the stem lysine residue (Schleifer and Kandler, 1972). PG subunits are polymerized into glycan chains by glycosyltransferase in penicillin binding proteins (PBPs) and by Shape, Elongation, Division, and Sporulation (SEDS)-family proteins, RodA and FtsW (Meeske et al., 2016). Adjoining glycan chains are then crosslinked via transpeptidation, where the transpeptidase activity of PBPs forms a peptide bond between the L-alanine crossbridge and the D-alanine residue at the fourth position of the pentapeptide stem. During crosslinking, the terminal D-alanine of the pentapeptide stem is removed. Although the general PG subunit is similar for most Gram-positive bacteria, the amino acid sequence of the crossbridge can vary. The crossbridge of *E. faecalis* consists of two L-alanines while the crossbridge of *Enterococcus faecium*, *Streptococcus pneumoniae* and *S. aureus* are composed of a single D-aspartate, L-alanine-L-alanine or L-serine-L-alanine and five glycines, respectively (Bellais et al., 2006; Fiser et al., 2003; Schneider et al., 2004). The differences in crossbridge length in turn affect the extent of cell wall crosslinking whereby a longer crossbridge is associated with a more crosslinked cell wall and vice versa (Kim et al., 2014). For example, the percentage of crosslinking of *S. aureus* ranges from 74 – 92% (Vollmer and Seligman, 2010), whereas the percentage crosslinking in *E. faecalis* and *S. pneumoniae* is approximately 48% and 35%, respectively (Bui et al., 2012; Yang et al., 2017).

While the process of pilus assembly, sorting machinery, and cell wall synthesis are interlinked and well-studied as individual processes, little is known about the interplay between these mechanisms. Moreover, despite differences in cell wall structure among Gram positive bacteria, the main paradigm for cell wall protein surface deposition is based on studies carried out in *S. aureus*. Here, we sought to bridge the gap between cell wall protein anchoring and cell wall synthesis by investigating the spatial and temporal distribution of surface-exposed pilus in *E. faecalis*. Since *E. faecalis* sortases are focally enriched at the cell septum (Kline et al., 2009), which coincides with the site of new cell wall synthesis, we hypothesized that new Ebp would be also emerge focally at the division septum. To our surprise, we found that Ebp emerge and localize at the cell periphery, predominantly saturating one hemisphere of the diplococcus, and are excluded from the cell septum. Data in this manuscript support a model in which *E. faecalis* SrtA substrates do not become incorporated into the cell wall not at sites of new wall synthesis, but rather at sites of uncrosslinked older PG at the cell periphery.

## Methods

### Bacterial culture and strains

Bacterial strains and plasmids used in this study are listed in **Table S1**. Unless otherwise stated, *E. faecalis* strains were streaked onto brain heart infusion (BHI) agar from 25% glycerol stocks stored at −80°C and grown overnight at 37 °C. Single colonies were then inoculated into brain heart infusion broth (BHI broth; BD Difco, USA) and grown overnight statically for 16 to 18 hours at 37 °C. To obtain cells at mid log phase, the overnight cultures were subcultured at 1:10 dilution into fresh BHI media and grown to OD_600_ 0.5 ± 0.05. Cells were then normalized to OD_600_ 0.5 in 1 mL 0.1 M phosphate buffer (PB) for subsequent use unless otherwise stated. Where appropriate, antibiotics were added at the following concentrations: tetracycline (Tet), 15 μg/ml; Kanamycin (Kan), 500 μg/ml.

### Genetic manipulation

*E. faecalis* OG1RFΔ*srtA* was transformed with plasmid pGCP123::P*srtA srtA-2L-mCherry* in which SrtA-mCherry fusion is expressed on a plasmid and transcribed from the native *srtA* promoter. The wild-type *srtA* native promoter was amplified using primers KpnI-PsrtA-F (5’-ATCCGGTACCGCTTGTTTCTTTTACTTTAAAATTCCA-3’) and XhoI-PsrtA-R (5’-AAGCCTCGAGATTCTCCCTCCTTTTAATGT-3’) and the *srtA* gene was amplified using primers XhoI-SrtA-F (5’-GAATCTCGAGATGCGCCCAAAAGAGAAAAA-3’) and EcoRI-SrtA-R (5’-ATCCGAATTCAGCCACCCAATCGGCTAA3’), using *E. faecalis* OG1RF as template. mCherry was amplified using primers EcoRI-SrtA-R (5’-ATCCGAATTCATGGTGAGCAAGGGC-3’) and NotI-STOP-XbaI-BamHI-mCherry-R (5’-AATCGCGGCCGCCTATCTAGAGGATCCCTTGTACAGCTCGTCCAT-3’). These three PCR products were ligated together using primer embedded restriction sites XhoI and EcoRI respectively. The resulting fusion product was cloned into PGCP123 (Nielsen et al., 2012) using primer embedded restriction sites KpnI and NotI. The expression and stability of the fusion was verified with anti-mCherry (Invitrogen) and anti-SrtA (SABio, Singapore) immunoblotting of whole-cell *E. faecalis* lysates.

### Immunofluorescence microscopy

Immunolabelling of cells was performed as described in (Kandaswamy et al., 2013) with modifications. Mid log phase cells were normalized to OD_600_ 0.5 and washed once in 0.01M low salt phosphate buffer (PB). Cells were then fixed in 4% (wt/vol) paraformaldehyde (PFA) and incubated for 20 min at room temperature (RT). After fixation, cells were either treated with 10 mg/mL lysozyme for 37°C for 1 hour to remove the cell wall for labelling of membrane-bound Ebp, or lysozyme untreated for labelling of surface-exposed Ebp. After washing once with PB, cells were incubated in 2% (wt/vol) bovine serum albumin (BSA) in PB for 20 minutes at RT. For Ebp staining, cells were incubated with guinea pig anti-EbpC serum or rabbit anti-EbpA serum (Afonina et al., 2018) at 1:500 dilution in PB-2% BSA for 1 hour at RT. Next, cells were washed once in PB and incubated with fluorescent-conjugated secondary antibody at 1:500 dilution in PB-2% BSA for 1 hour (Alexa Fluor 405/488/568– goat anti-guinea pig antibody for EbpC and goat anti-rabbit antibody for EbpA [Invitrogen, Inc., USA]). Cells were then washed once in PB and resuspended in 1 mL PB. Before sample mounting, microscope glass slides were washed once in filtered 70% ethanol followed by filtered ultrapure water (18.2 ohm) and dried. Hydrophobic wells measuring 1.25 cm in diameter were drawn on the glass slide using a PAP pen (Sigma Aldrich, Singapore). 20 µL of cell suspension was spotted onto each well and allowed to dry in a 60 °C oven. Samples were covered with 5 µL of mounting media (Vectashield®, USA) and sealed with glass coverslips. Widefield microscopy was performed using an inverted epi-fluorescence microscope (Zeiss Axio observer Z1, Germany) fitted with a Plan-Neofluar 100x/1.3 oil Ph3 objective lens using ZEN 2 (blue edition) software. The images were acquired using AF568/cy3 filter cube sets fitted with a 530–580 nm bandpass excitation filter and a 585 nm-long pass barrier filter. For unbiased image analysis, exposure times were fixed for all experiments. Images were processed using FIJI (Schindelin et al., 2012) and Adobe Photoshop CS5.1.

### Super-resolution structured illumination microscopy (SIM)

All SIM imaging were performed using an alpha Plan-Apochromat 100x/1.46 oil DICIII objective lens and pco.edge sCMOS camera fitted onto an Elyra PS.1 microscope (Zeiss). Laser wavelengths of 561, 488 and 405 nm at 20% power were used to excite red, green and blue fluorescent probes respectively. Images were acquired using five grid rotations with 51 μm grating period and reconstructed using ZEN software (ZEN 2012 SP5 FP2, black edition).

### Quantitative analysis of Ebp fluorescence distribution on single cells

Fluorescence distribution of Ebp staining was quantified as described previously (Chilambi et al., 2020; Kandaswamy et al., 2013). Briefly, mid log phase cells (1.5 µm −2 µm in length) were first detected using a Matlab function, Projected System of Internal Coordinates from Interpolated Contours (PSICIC) (Guberman et al., 2008). Cell perimeters of each cell were traced and given an arbitrary unit from 0 to 100 where positions 0, 100 and 50 mark the poles of the cell while positions 25 and 75 mark the division septum of the cell. The orientation profiles of cells were manually arranged such that 0 always corresponds to the pole with lower fluorescence intensity. Fluorescence intensity values corresponding to each position on the cell were plotted to generate a fluorescence distribution pattern.

### Fluorescent D-amino acid (FDAA) labelling

Cells were subcultured 1:10 in 5 mL BHI and grown to early log phase (OD_600_ 0.25 ± 0.05). Cells were normalized to OD_600_ 0.25 in pre-warmed BHI and the FDAA, 7-hydroxycoumarin-3-carboxylic acid 3-amino–D-alanine (HADA) (Kuru et al., 2015), was added to a final concentration of 250 µM. Cells were then grown for either 5, 20, 40 or 120 minutes at 37 °C with agitation. To halt the labelling, cells were placed on ice and washed thrice in ice cold PB. Cells were then mounted onto clean glass slides before imaging by SIM. For FDAA sequential labelling, cells were labelled first with the green derivative of BODIPY-FL 3-amino-D-alanine (BADA) for 5 minutes or 40 minutes at 37 °C in BHI, washed once in ice cold PBS and then labelled with the red FDAA derivative, TAMRA 3-amino-D-alanine (TADA) for 5 minutes or 40 minutes at 37 °C in BHI. Cells were then washed once in ice cold PBS and incubated in BHI with HADA for 10 minutes or 20 minutes at 37 °C, washed once in PBS and mounted onto glass slides and imaged via SIM as described above.

### Time Lapse imaging

BHI agarose (1%) gel pads were prepared by sandwiching molten BHI agarose between two glass slides. Once solidified, 1 cm by 1 cm agarose gel squares were cut out. 5 µL of mid log phase cells, pre-labelled for Ebp were spotted and evenly spread across the agarose gel pad and sealed with a glass cover slip using paraffin wax. Images were then taken every 15 minutes at room temperature for 2.5 hour using an inverted epi-fluorescence microscope (Zeiss Axio observer Z1, Germany) fitted with a 100x/1.3 oil Ph3 objective lens.

### Triple Ebp chase labelling

*E. faecalis* OG1RF cells were grown in BHI to mid log phase and stained for EbpC via immunofluorescence first with a green fluorescent secondary antibody, Alexa Fluor 488 (Invitrogen, Singapore). After the first Ebp labelling, cells were washed and further grown in BHI for 1 hour at 37 °C, followed by staining for EbpC with a blue fluorescent secondary antibody, Alexa Fluor 405. Next, the cells were washed and again allowed to grow in BHI for 1 hour at 37 °C and similarly labelled for EbpC but with a red fluorescent secondary antibody, Alexa Fluor 568. After the final staining, cells were mounted and imaged via SIM.

### Determination of subbacteriostatic concentration of antibiotics

*E. faecalis c*ells were grown to OD 0.4 at 37 °C and subsequently transferred into a 96-well plate containing ramoplanin (0, 16 – 30 μg/mL) in BHI. Cells were incubated in ramoplanin at 37 °C for 2 hours and absorbance (600 nm) was read every 15 minutes using a Tecan M200 microtiter plate reader. The sub-bacteriostatic concentration of ramoplanin was determined as the lowest dose of antibiotics that showed a decrease in absorbance reading within the first hour of incubation.

### Ramoplanin Ebp-HADA chase experiment

*E. faecalis* cells were grown to mid log phase, followed by EbpC labelling using a green fluorescent secondary antibody, Alexa Fluor 488. After washing once in PB, cells were grown in the presence or absence of ramoplanin with 250 µM HADA in BHI at 37 °C for 1 hour. Cells were washed and again labelled for EbpC, instead with a red fluorescent secondary antibody, AlexaFlour 568. SIM imaging was performed as described above.

## Results

### Cell surface-exposed Ebp are septum exclusive and temporally deposited towards the pole

Sorting and cell surface-exposure of Enterococcus pili can facilitate its virulence during infection. We visualized the distribution of fluorescently labelled *E. faecalis* pili at three different growth phases, defined as 1) early division elongated monococci which have not yet undergone septation (1 µm −1.5 µm in length), 2) mid division diplococci which are undergoing elongation and septum constriction (1.5 µm −2 µm), and 3) late division cells consisting of two daughter cells just before separation (> 2 µm). During each growth phase, we observed EbpC labelling only at the cell hemispheres and at sites adjacent to the equatorial ring, which corresponds to the site of future cell division, but not at the septum, a pattern we refer to as “septum exclusive” (**Fig. 1A**). Furthermore, we observed that one hemisphere of the cell is always more saturated with Ebp than the other. The hemispherical saturation and septum exclusive localization patterns were seen in early and mid-division phase cells. However, in late division cells, we observed EbpC fully covering both hemispheres with two additional foci adjacent to the nascent septum, mirroring each other (**Fig. 1A**). The transition from asymmetric distribution of Ebp in the mid-division phase to symmetric Ebp distribution between the two hemispheres in late division phase hinted that the two foci adjacent to the septum may seed the eventual coverage of the entire hemisphere. To gain a more detailed characterization of Ebp distribution, we quantified the localization pattern of EbpC by plotting the Ebp fluorescence intensity against the cell perimeter in mid-division phase cells and observed that the fluorescence intensity is lowest at positions that correspond to the septum (**Fig. 1B**). The low fluorescence intensity at the septum is consistent with our observation that the localization of cell wall associated Ebp is septum exclusive. Moreover, the fluorescence intensity is highest from positions 38 to 63, which correspond to the beginning and end of one hemisphere of the cell, again consistent with our observation that one side of the cell is more saturated with Ebp than the other.

**Fig. 1.**
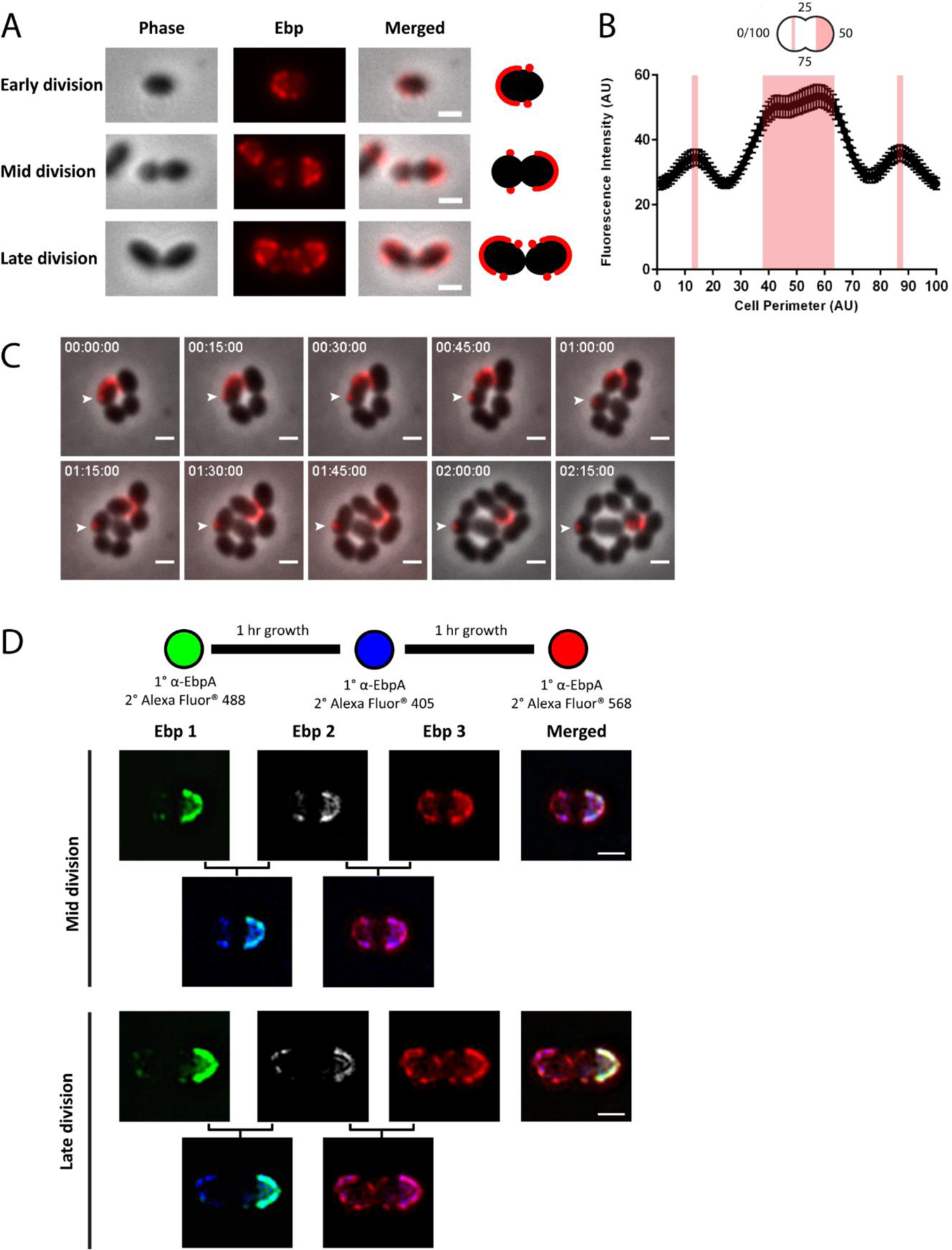
Distribution of surface-exposed Ebp in *E. faecalis*. **(A)** Representative immunofluorescence (IF) labelling of surface-exposed Ebp in *E. faecalis* at early, mid and late division. Scale bar, 1 µm**. (B)** Quantification of Ebp fluorescence intensity along the cell periphery (0-100) of mid division cells starting from the cell pole where 25 and 75 correspond to the cell septum (n=92). The shaded areas on the graph correspond to one hemisphere and two foci adjacent to the equatorial ring with peak fluorescence intensity. **(C)** Time-lapse of live *E. faecalis* cells pre-labelled for Ebp, mounted on a BHI agarose pad and imaged every 15 minutes. **(D)** Chase labelling of Ebp via immunofluorescence at 1 hour growth intervals using green, blue and red fluorescent conjugated secondary antibodies as shown in the schematic. Images on the second row are merged imaged of either green and blue or blue and red. Representative mid and late division phase cells are shown. Scale bar: 1 µm

Two smaller peaks were observed at positions 13 and 87, which correspond to the two foci located adjacent to the septum on the other hemisphere of the cell (**Fig. 1B**). Because the maximum fluorescence intensity of these two foci is lower than the other hemisphere, we hypothesized that these two foci represent newly emerged Ebp and that additional Ebp would continue to emerge to eventually complete the hemispherical coverage. Overall, these findings indicate that pilus localization is septum exclusive and that Ebp decoration predominates at one hemisphere of the cell.

Because we observed that Ebp become surface-exposed (and therefore available for binding by extracellular antibodies for immunofluorescence) and saturates one cell hemisphere before the other, we hypothesized two possible scenarios to describe how one hemisphere becomes fully piliated. In the first scenario, the two foci adjacent to the septum remain immobile and newer pili emerge next to them, proceeding toward the pole, where they eventually converge to cover the hemisphere. In the second scenario, the two foci migrate towards the pole while new Ebp continue to emerge behind the migrating foci from the site adjacent to the septum. To determine which of these two hypotheses could explain temporal pilus deposition on the cell wall, we performed time-lapse fluorescence microscopy on live EbpC-labelled cells. We traced the original fluorescent Ebp foci on the cell and observed that as the cell elongates and divides, the single focus does not move along the cell periphery. Instead, the stained EbpC focus remains where it was originally labelled, while new cell material is synthesized at the septum as the cell divides (**Fig. 1C**).

This lack of focus movement suggested that once deposited on the cell wall, Ebp are immobile and that newer Ebp are likely deposited towards the cell pole to achieve their hemispherical localization pattern. To track newly emerged Ebp that are predicted to emerge towards the cell pole, we performed an Ebp pulse chase labelling experiment where Ebp were labelled three consecutive times via immunofluorescence using green, blue, and red fluorescently labelled secondary antibodies sequentially with a washout step and 1 hour of growth between each labelling. We imaged the cells using structured illumination microscopy (SIM) and observed that Ebp were labelled in a sequential manner towards the pole (**Fig 1D**). In the hemisphere where the two Ebp foci were initially labelled in green, we observed blue-labelled Ebp overlap with the older green-labelled Ebp, in addition to new blue-label localizing to segments adjacent to the green foci. On the same hemisphere, red-labelled Ebp, representing the newest Ebp deposited on the cell surface, overlapped with both green and blue and covered the entire cell hemisphere. To test if other cell wall anchored substrates were exposed in a similar manner, we performed similar chase labelling experiment on aggregation substance (AS) and observed similar sequential deposition of AS on the cell (**Fig. S1**).These data were consistent with our observation on live cells that Ebp deposited on the cell wall are fixed in space and that newer Ebp are deposited in a temporal manner towards the cell pole where, over time, newer Ebp are exposed closer to the pole while older Ebp remain closer to the equatorial rings until eventually, the whole hemisphere is saturated with Ebp. These data also hinted that Ebp deposition and cell wall synthesis are coordinated where new Ebp may appear together with newly synthesized cell wall.

### New cell wall is synthesized at the septum and is driven towards the cell hemisphere as the cell elongates

The current paradigm in Gram-positive bacteria, largely defined in *S. aureus,* is that SrtA substrates are anchored to the cell wall via lipid II, a cell wall precursor (Ton-That and Schneewind, 1999; Ton-That et al., 1997). Similarly, we postulated that, in *E. faecalis*, sortase-anchored Ebp is attached to lipid II and becomes surface-exposed as this lipid II-substrate intermediate is incorporated into the growing PG cell wall. To test this, we first characterized *E. faecalis* cell wall synthesis dynamics using fluorescent D-amino acid (FDAA) probes. FDAAs are fluorescently-modified D-amino acids that are actively incorporated into the cell wall by D,D- and L,D-transpeptidases (Hsu et al., 2017; Kuru et al., 2015). We incubated mid log phase cells with HADA, a blue-emitting FDAA, for 5, 20, 40 or 120 minutes and visualized the staining patterns using SIM (**Fig. 2A**). After 5 minutes of HADA exposure, we observed cell wall labelling exclusively at the septum, indicating that new cell wall is synthesized at mid cell, as expected and reported for other related species (Boersma et al., 2015; Hsu et al., 2019). After 20 minutes of HADA exposure, we observed a distinctive “cross-like” localization pattern where cell wall was stained from the equatorial ring to the septum and unlabelled at both hemispheres (**Fig. 2A**). The cross-like localization pattern supports that new cell wall is synthesized at the mid-cell. By contrast, after 40 minutes of labelling, most cells were stained at the septum and one side of the hemisphere while the other hemisphere remained unlabelled (**Fig. 2A**). This differential distribution of PG staining is consistent with one half of each diplococcus being more mature than the other. After 120 minutes of HADA exposure, most cells were uniformly labelled, indicating that they have gone through at least 2 replication cycles (**Fig. 2A**). The cell wall labelling patterns observed after HADA labelling for different durations led us to postulate that PG incorporation in *E. faecalis* is coordinated asymmetrically. We proposed that new PG is always synthesized at midcell and older PG is consistently, constantly, and evenly pushed out by newer cell wall with little or no intercalation of new and old cell wall material.

**Fig. 2.**
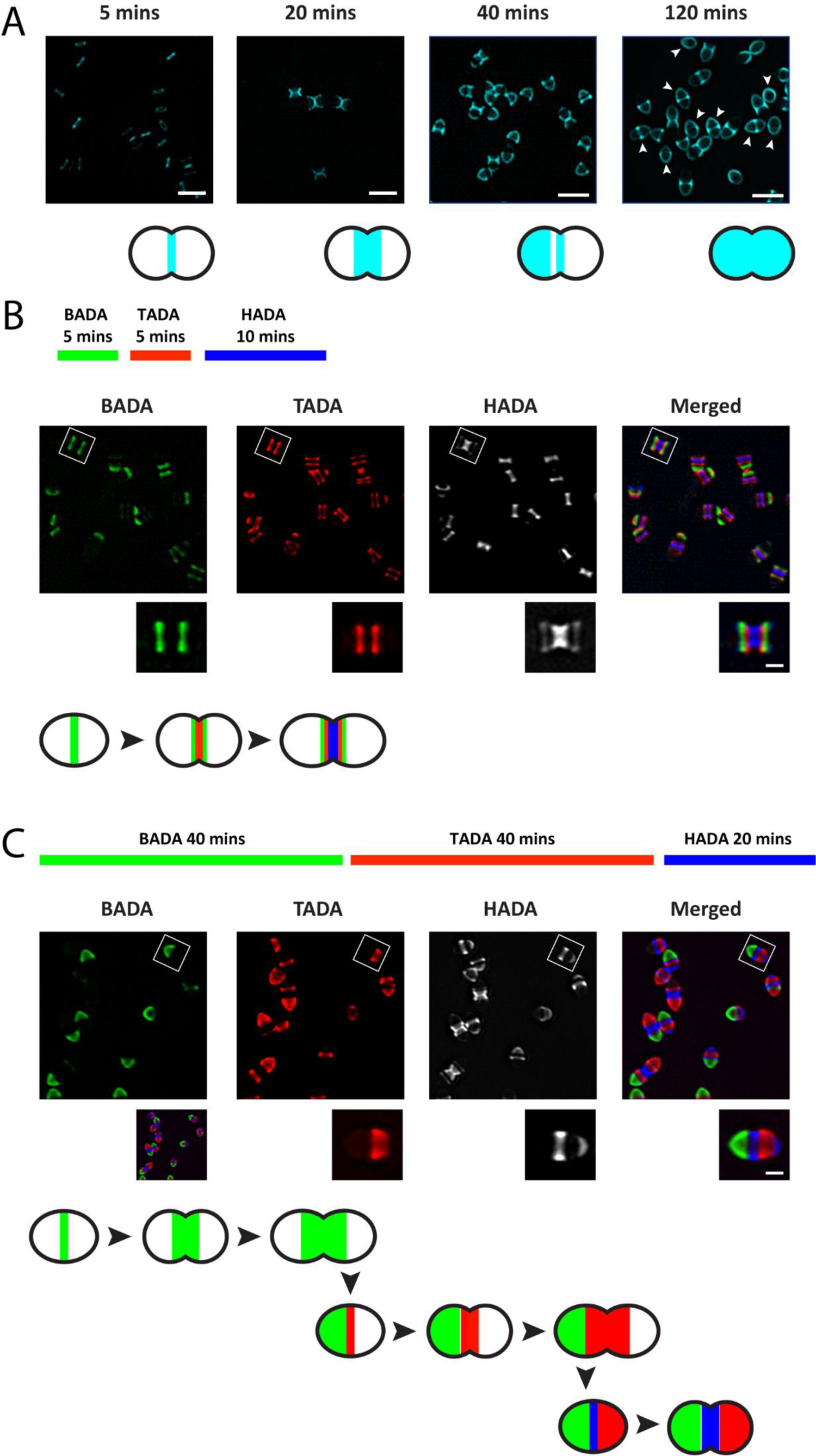
Fluorescent D-amino acid (FDAA) labeling of *E. faecalis* cell wall throughout the cell cycle. **(A)** Representative images of HCC-amino-D-alanine (HADA) labelled *E. faecalis* cell wall, pseudocolored cyan, at various time points. Cells were grown to exponential phase and labelled with HADA for 5, 20, 40 and 120 minutes. White arrows indicate cells that are fully labelled with HADA. Cells were imaged by structured illumination microscopy (SIM). HADA has been pseudo-colored cyan for ease of visualization. Scale bar: 1 µm. **(B & C)** Representative images after sequential labelling of *E. faecalis* cell wall at short **(B)** and long **(C)** pulses and imaged by SIM. In **(B)**, cells were labelled with BADA (5 minutes), TADA (5 minutes) and HADA (10 minutes). In **(C)**, cells were labelled with BADA (40 minutes), TADA (40 minutes) and HADA (20 minutes). Scale bar: 0.5 µm. Schematics of representative cells are shown at the bottom of each panel.

To spatially distinguish between old and new cell wall, and to determine the temporal distribution of PG, we next performed a sequential labelling of new PG using three different colored FDAAs. A series of short pulses was carried out, in which we first incubated the cells for 5 minutes with a green FDAA, BODIPY-FL 3-amino-d-alanine (BADA) followed by 10 minutes with a red FDAA, TAMRA 3-amino-D-alanine (TADA) and lastly, 10 minutes with HADA (**Fig. 2B**). The three stains appeared as distinct fluorescent bands parallel to each other on the cell with minimal overlap, suggesting immobility of the cell wall PG after it is incorporated. BADA and TADA fluorescent bands appeared narrower than that of HADA’s, suggesting splitting of these fluorescent bands at midcell as newer PG is synthesized and incorporated at the septum. Furthermore, BADA labelled PG formed the outermost bands, flanking both TADA and HADA labelled PG with TADA sandwiched between BADA and HADA (**Fig. 2B**). This chronological labelling pattern supports our hypothesis that PG at midcell is pushed away from the septum as the cell elongates and divides. We further demonstrated the ability to visualize the older hemisphere of the cell by extending the serial FDAA labelling of BADA, TADA and HADA to 40 minutes, 40 minutes, and 20 minutes, respectively (**Fig. 2C**). We chose 40 minutes for the first two pulses because we observed single hemispherical PG staining for this staining duration and if our hypothesis that one hemisphere of a diplococcus is always older than other is true, we predicted that the older hemisphere of the cell will be labelled green, the other hemisphere labelled red, and the middle of the cell labelled blue. Consistent with this prediction, BADA labelling was seen predominantly at one hemisphere while TADA labelled the other cell hemisphere, and HADA staining was observed in the middle of the cell (**Fig. 2C**). These data supported our hypothesis that the cell wall of one hemisphere is always older than the other. Our findings demonstrate that cell wall synthesis in *E. faecalis* occurs strictly at the septum and there is minimal overlap between the old and new cell wall.

### Ebp does not co-localize with newly synthesized cell wall

Ebp is a surface-exposed virulence factor that is covalently attached onto the cell wall by SrtA (Nielsen et al., 2013). Based largely on studies in *S. aureus* (Perry et al., 2002; Ton-That and Schneewind, 1999; Ton-That et al., 1997), SrtA is thought to anchor its substrates onto the growing cell wall via linkage to a lipid II precursor. Therefore, we postulated that Ebp is exposed to the cell surface together with newly synthesized cell wall at the septum, via a similar interaction with lipid II precursors. To test if Ebp exposure coincides with the new cell wall, we performed co-localization studies by co-staining both the cell wall and pili (**Fig. 3AB**). We performed both short and long incubations with HADA to determine if there were any differences in co-localization patterns. Surprisingly, Ebp did not co-localize with the newly synthesized labelled cell wall regardless of the length of HADA exposure. Instead, we saw Ebp staining where HADA labelling was absent. Ebp labelling also appeared more saturated at the older hemisphere. Our results illustrate that a mature cell wall may be required before *Enterococcus* pilus can be exposed on the surface and accessible to antibody labelling.

To test if PG synthesis precedes Ebp deposition, we performed a HADA-Ebp chase experiment. The experimental design was similar to the above assay, but included additional steps where cells were allowed to grow in BHI for an hour (in the absence of any stain) and then stained again for Ebp with a green fluorescent secondary antibody. When cells were exposed to HADA for 1 hour, costained with Ebp, allowed to grow, and then stained a second time for Ebp, we saw that newly labelled Ebp overlapped with the pre-labelled cell wall (**Fig. 3C**). This result confirmed our hypothesis that the cell wall must undergo cell growth-associated wall maturation before Ebp is anchored onto it. We propose a model associating the process of Ebp exposure and new cell wall synthesis during cell division

**Fig. 3.**
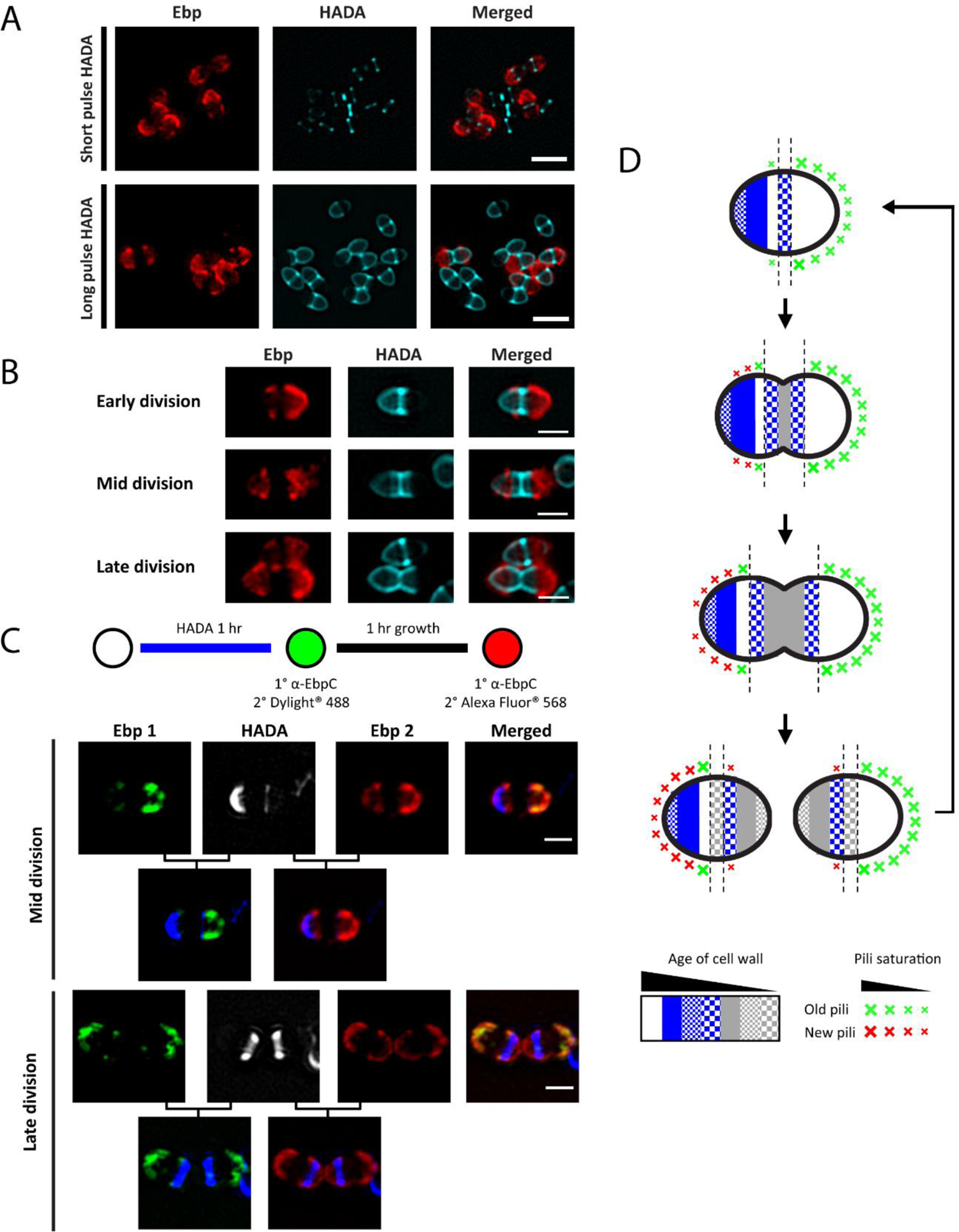
Co-staining of peptidoglycan and Ebp in *E. faecalis*. **(A & B)** Exponential growing cells were labelled with HADA for short pulse (5 minutes) or long (120 minutes), stained for Ebp via immunofluorescence and imaged via SIM. Scale bar = 2 µm. **(B)**, Representative image of cells at early, mid and late division phases are shown. Scale bar = 1 µm. **(C)** HADA staining followed by chase labelling of Ebp. Exponentially growing cells were harvested, stained with HADA for 1 hour followed by labelling of EbpC via immunofluorescence first with a green and then a red secondary antibody after 1 hour growth in BHI. Images were taken via SIM and two representative images are shown. Scale bar = 1 µm. **(D)** Proposed model of pili exposure in relation to cell wall synthesis during cell division. Cells are segmented by age of the cell wall as determined in Fig. 2, where white is the oldest followed by blue (solid), blue (small checker board), blue (large checker board), grey (solid), grey (small checker board) and grey (large checker board). Sites of old exposed pili are labelled green while sites of newly exposed pili are labelled red. Initially, pili are saturated at the older hemisphere of the cell and also appear as two foci adjacent to the septum on the younger hemisphere. As the cell elongates and divides, new pili on the younger hemisphere are exposed sequentially towards the cell pole in a cell wall age dependent manner. The younger cell hemisphere is eventually fully saturated with surface-exposed pili as the cell wall matures.

### Pili cell surface exposure location is independent of focally enriched septal SrtA

Previously, we showed that SrtA was enriched at the *E. faecalis* cell septum (Kandaswamy et al., 2013; Kline et al., 2009). Yet in this study, we show that surface exposed Ebp, a SrtA substrate, is localized away from the cell septum and appears in a cell wall age-dependent manner at the cell hemispheres. We postulated that, even though SrtA is enriched at the septum, peripheral SrtA may anchor Ebp to the hemispherical cell wall. To examine SrtA localization relative to Ebp, we labelled surface exposed Ebp in a *srtA*-mCherry *E. faecalis* strain, enabling visualization of membrane-anchored SrtA in cells with intact cell wall (compared to immunolabeling of SrtA, which requires lysosome removal of the cell wall to render the membrane protein antibody accessible). Consistent with previous studies, we saw SrtA focally enriched at the septum and at puncta along the cell periphery. Ebp localization was coincident with peripheral SrtA puncta, rather than septal SrtA in all division phases (**Fig. 4A**). This observation supports our data that Ebp becomes surface exposed at the cell hemispheres, and may be anchored to the cell wall by peripheral SrtA. Polymerization of Ebp by SrtC occurs on the cell membrane, where membrane-bound polymerized pili are subsequently anchored it to the cell wall by SrtA (Nielsen et al., 2013). To investigate if sites of SrtA-dependent cell wall anchoring are only found at the cell hemispheres, we labelled Ebp in SrtA mutants, in which Ebp remains membrane-bound and is not attached to the cell wall. We predicted that if Ebp are attached to the cell wall at the periphery, they might similarly accumulate in the membrane in a peripheral and hemispherical pattern in a Δ*srtA* strain. We first examined Ebp by immunolabelling of EbpC prior to lysozyme treatment for cell wall removal. Contrary to our expectation that we would not see any surface exposed Ebp in Δ*srtA*, we instead saw hemispherical labelling of Ebp on the cell surface, similar to WT cells (**Fig. 4B**), which we presumed to be anchored in the membrane. Consistent with previous reports, we verified by immunoblot that the surface exposed Ebp in Δ*srtA* are indeed membrane bound (Nielsen et al., 2013) (**Fig. S2**). Together, the immunolabelling and immunoblot results tell us that despite polymerized pili being membrane bound when SrtA is absent, these pili can protrude out of the cell wall and be detected at the surface of the cell hemispheres.

**Fig. 4.**
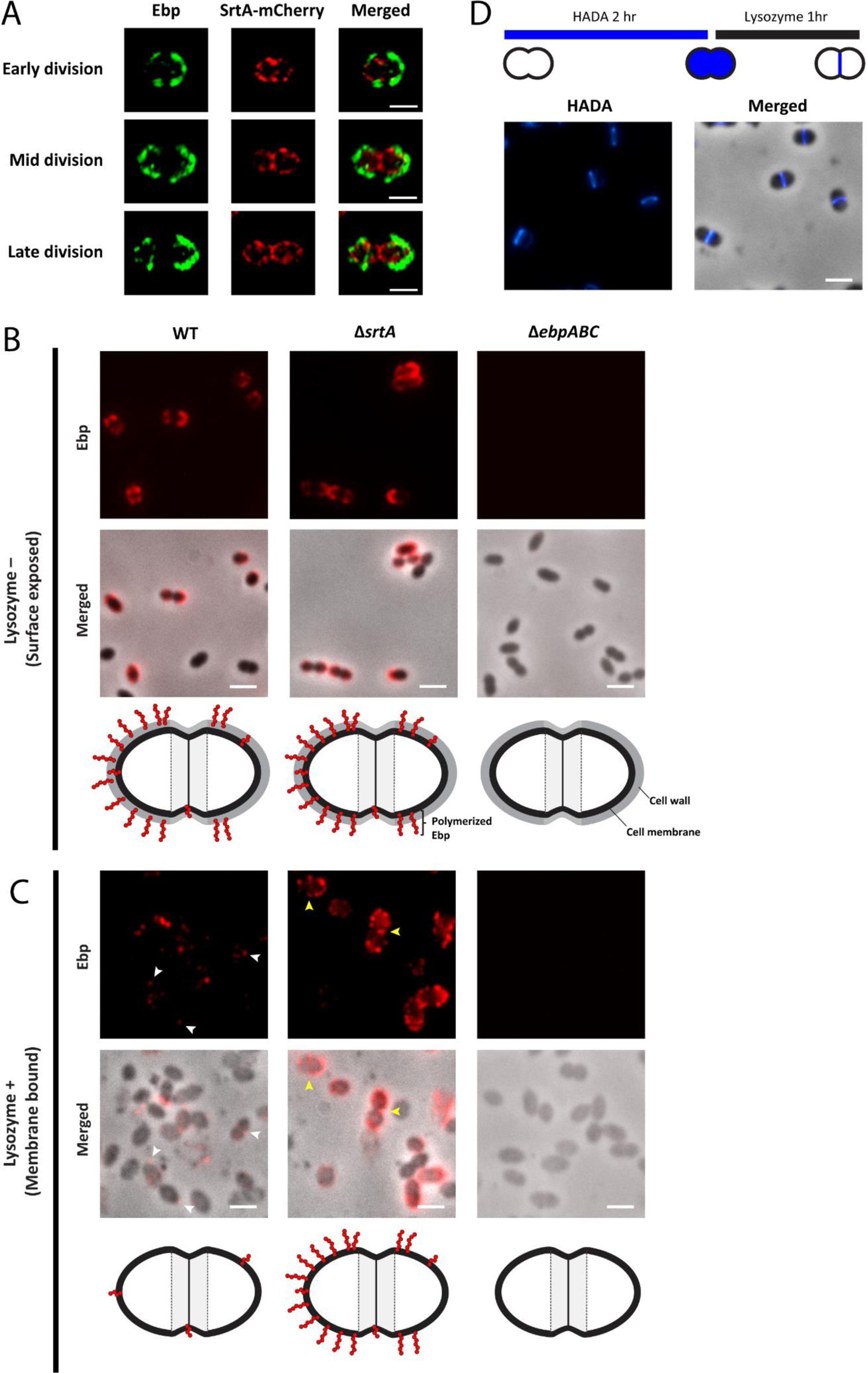
SrtA and Ebp co-labelling and distribution of cell wall- and membrane-bound pili in *E. faecalis*. **(A)** Representative SrtA-mCherry expressing cells, co-labeled with Ebp and imaged by SIM. Scale bar: 1 µm **(B)** Lysozyme untreated cells labelled for Ebp and imaged via epifluorescence and phase contrast microscopy. Scale bar: 2 µm **(C)** Lysozyme treated cells labelled for Ebp and imaged via epifluorescence and phase contrast microscopy. White arrows indicate stained Ebp on the surface of *E. faecalis* membranes in WT. Yellow arrows point to Ebp localized septally in Δ*srtA*. Scale bar: 2 µm. Cartoon images on last panel of **(B)** and **(C)** depicts proposed Ebp positions on the respective strains. **(D)** Images of cells lysozyme treated for 1 after pre-labelling the cell wall with HADA for 2 hours. Scale bar: 2 µm.

After cell wall removal by lysozyme treatment, we expected to see no Ebp labelling on the protoplast of wild type (WT) cells as Ebp anchored to the cell wall would have been removed. Strikingly, even though we no longer saw hemispherically localized Ebp, we observed cells decorated with Ebp foci at random locations on the protoplasts (**Fig. 4C**). No fluorescence was detected in Δ*ebpABC* protoplasts, indicating that the anti-EbpC antibody was specific (**Fig. 4C**). To ensure that the Ebp on the protoplast was not a result of incomplete removal of the cell wall, we performed the same lysozyme treatment on cells pre-labelled with HADA for 2 hours and observed complete removal of the peripheral cell wall (**Fig. 4D**, compared to Fig. 2A where HADA labelled cells were imaged in the absence of lysozyme treatment). We hypothesized that the Ebp foci observed on WT protoplasts could be sites of polymerized and membrane anchored pili prior to cell wall incorporation by SrtA. The presence of Ebp localized at the cell septum in WT cells suggests that Ebp might be polymerized at the septum and not exposed to the surface yet. Moreover, in lysozyme treated Δ*srtA*, apart from hemispherical labelled membrane bound Ebp, Ebp foci could also be found at the septal region of some cells (**Fig. 4C**). The presence of Ebp septal labelling on Δ*srtA* protoplast supports our speculation that newly polymerized pili may be buried under the septal cell wall and not yet surface exposed. Coupled with the earlier observation that pilus exposure on the cell surface is related to the age of the cell wall, we speculate that the similar hemispherical localization pattern between surface-exposed membrane-bound Ebp and cell wall anchored Ebp correlates with cell wall maturity, where older cell wall may be more porous due to growth-associated cell wall maturation (Pasquina-Lemonche et al., 2020), thus allowing Ebp surface exposure.

### New pilus deposition at the cell hemisphere is independent of new cell wall synthesis

The synthesis and assembly of the *E. faecalis* cell wall components are crucial in promoting its virulence, however the coordination between these different processes remain un-elucidated. Polymerized pili are anchored onto the cell wall by SrtA (Nielsen et al., 2013). The sorting machinery was characterized in a *S. aureus* model whereby SrtA catalyzes the formation of a substrate-lipid II intermediate which is eventually incorporated into the cell wall via transglycosylation and transpeptidation reactions (Perry et al., 2002). Our observations that new cell surface exposure of Ebp and formation of the Enterococcal cell wall do not occur simultaneously led us to postulate that Ebp may bypass precursor lipid II and instead be directly anchored onto the uncrosslinked cell wall by SrtA. The *E. faecalis* cell wall is about 50% crosslinked compared to 85% in *S. aureus* (Kim et al., 2015; Yang et al., 2017). This lack of complete cell wall cross-linking suggests that there is an abundance of free L-Ala-L-Ala crossbridges in *E. faecalis* for anchoring of cell surface proteins. When *E. faecalis* was stained with BODIPY™ FL vancomycin (Vanc-FL), a fluorescently labelled antibiotic that targets the D-Ala-D-Ala residues in the cell wall (**Fig. 5A**), we saw enrichment of fluorescence at the septum and staining at the cell periphery (**Fig. 5B**). The intense fluorescence at the cell septum is indicative of the D-Ala-D-Ala on lipid II cell wall precursor, while the staining at the cell periphery indicates the D-Ala-D-Ala on uncrosslinked cell wall. To test if Ebp deposition is dependent on cell wall synthesis, and hence the availability of lipid II precursors, we treated cells with ramoplanin to inhibit cell wall synthesis. Ramoplanin primarily targets lipid II precursors, sequestering them, and rendering them unavailable for transglycosylation (**Fig. 5A**), and hence inhibiting new cell wall synthesis (Fang et al., 2006). We monitored growth in increasing concentrations of ramoplanin and determined the sub-bacteriostatic concentration to be 26 µg/mL in *E. faecalis* (**Fig. 5C**). To verify that cell wall synthesis was inhibited at this concentration, we grew ramoplanin-treated cells together with HADA. After 1 hour of incubation, no newly synthesised cell wall labelling was observed in ramoplanin-treated cells, confirming that cell wall synthesis was inhibited (**Fig 5D**). To determine if new Ebp was deposited onto the cell surface when cell wall synthesis was inhibited, we performed the HADA-Ebp chase experiment in the presence and absence of ramoplanin. Interestingly, at sub-bacteriostatic concentrations of ramoplanin, we saw new Ebp deposition at the cell hemisphere despite cell wall synthesis inhibition (**Fig. 5E**). To eliminate the possibility that new Ebp in ramoplanin treated cells were membrane-bound, we performed western blot to examine presence of the three subunits of Ebp in both the cell wall and protoplast fractions of both ramoplanin treated and untreated cells. However, the high molecular weight (HMW) ladder, representing polymerized pili, visible on the Ebp immunoblots were present only in the cell wall fraction in ramoplanin treated (**Fig. S3**), suggesting that the new Ebp are being incorporated into the cell wall via uncrosslinked PG away from the septum. Accordingly, when we stained ramoplanin treated cells with Vanc-FL, we observed homogeneous cell wall staining throughout the cell and lack of fluorescence enrichment at the septum, indicative of abundance of uncrosslinked PG in the cell and inhibition of new cell wall synthesis respectively (**Fig. 5B**). In contrast to the current paradigm for sortase substrate incorporation into the cell wall, our data suggest that in *E. faecalis* Ebp can be anchored onto uncrosslinked cell wall, which facilitates exposure to the cell exterior. We propose a new paradigm for *E. faecalis* Ebp deposition where SrtA mediated Ebp attachment of pili bypasses the need for lipid II-substrate intermediates and directly anchors pili onto uncrosslinked PG.

**Fig. 5.**
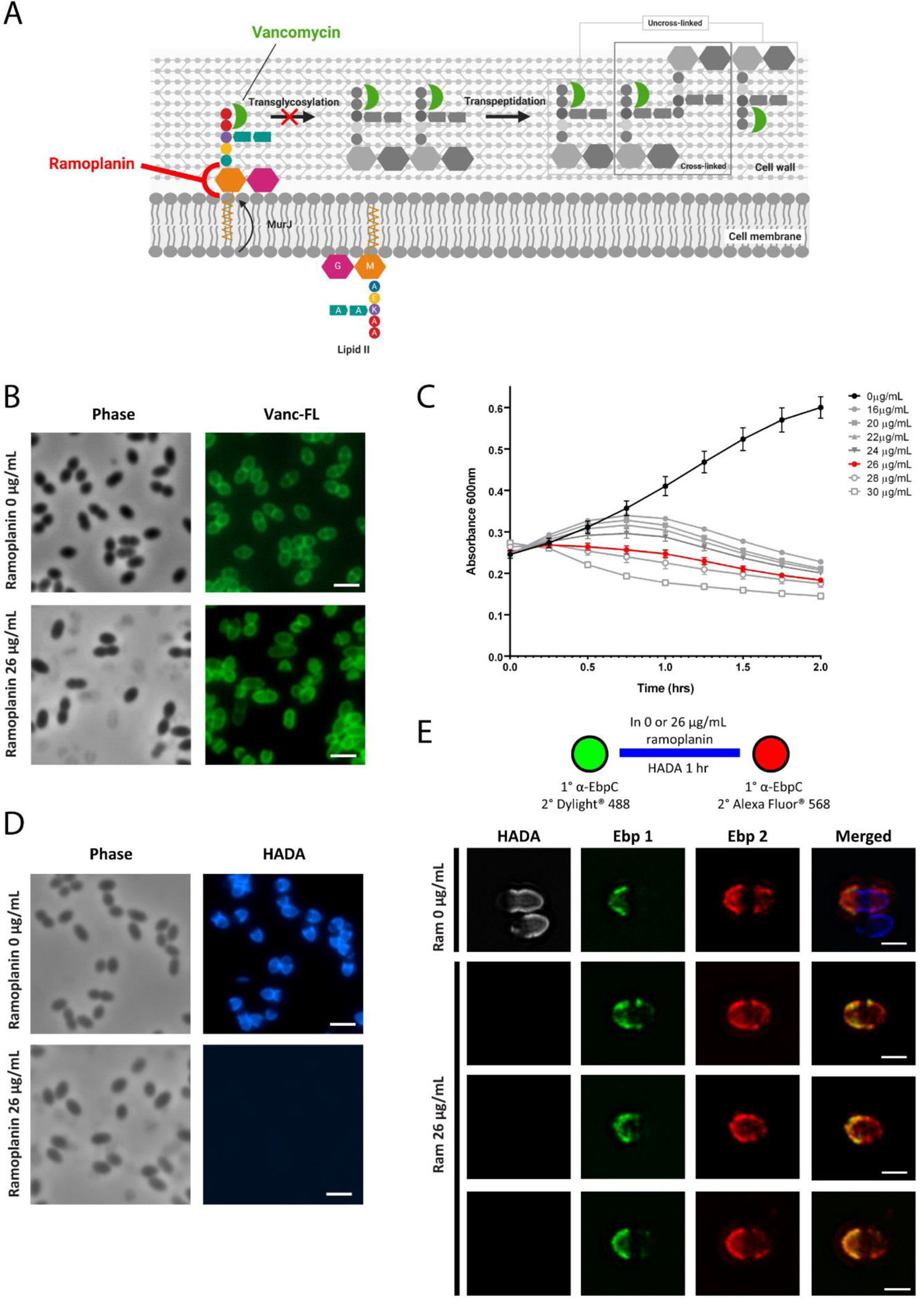
Cell wall synthesis inhibition by ramoplanin and its effect on pilus deposition. **(A)** Schematic showing target sites of cell wall inhibiting drugs, ramoplanin and vancomycin. **(B)** Cells treated or untreated with ramoplanin were stained with BODIPY® FL vancomycin and imaged. Scale bar: 2 µm **(C)** Exponentially growing bacteria were treated with increasing concentrations of ramoplanin for 2 hours at 37°C. Growth was measured at an absorbance of 600nm. **(D)** HADA labelling of bacteria treated in the presence or absence of 26 μg/mL ramoplanin for 1 hour. Scale bar: 2 µm **(E)** Exponentially growing cells were harvested and first labeled for Ebp in green via immunofluorescence. Cells were then allowed to grow in the presence or absence of ramoplanin with HADA for 1 hour. A second Ebp labelling was performed via immunofluorescence and imaged by SIM. Scale bar: 1 µm

**Fig 5.**
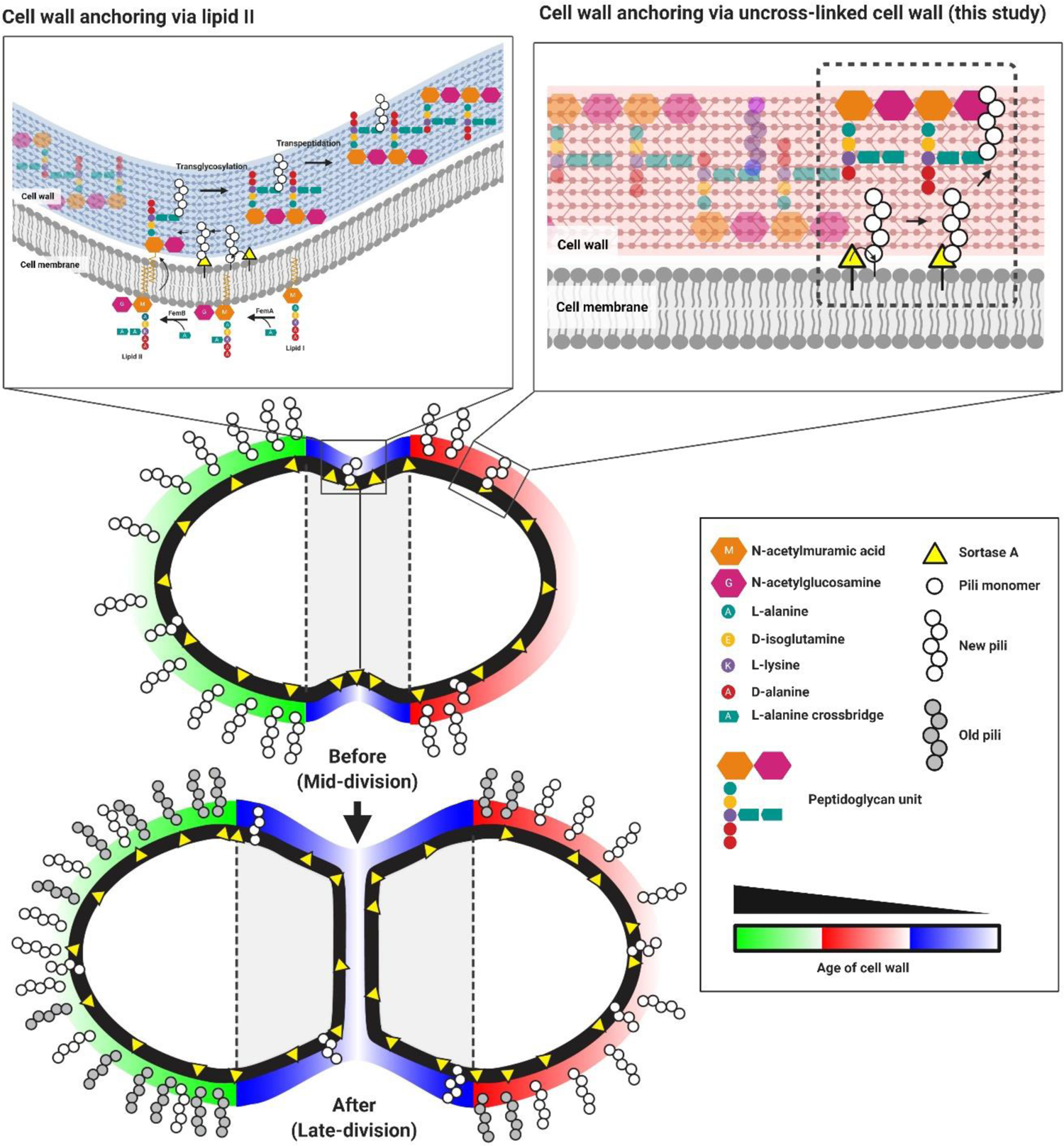
Spatial-temporal models of *E. faecalis* pili cell wall anchoring and pili deposition. Polymerized pili can either be anchored onto the cell wall via lipid II at the cell septum or uncrosslinked cell wall at the cell periphery. Surface-exposed pili are localized asymmetrically where the older hemisphere is more saturated with pili. Over time, the cell elongates and newer pili are deposited in a chronological manner towards the pole, eventually populating the whole hemisphere.

## Discussion

Spatial localization of cell wall-anchored surface proteins has been studied in various species of Gram-positive cocci. In *S. aureus* and *Streptococcus pyogenes*, cell-wall associated proteins are localized at the septal region or at the poles depending on the presence or absence of a conserved YSIRK motif within the substrates signal sequence, respectively (Carlsson et al., 2006; Raz et al., 2012). In *S. pneumoniae,* sortase-assembled pili are distributed in a non-homogenous manner, focally localized at the cell surface at discrete puncta (Fälker et al., 2008). In *E. faecium*, two distinct types of sortase-assembled pili are expressed, namely PilA and PilB. PilA are evenly distributed around the cell, while polymerized PilB are observed at the older pole at exponential phase (Hendrickx et al., 2008; Hendrickx et al., 2010). Hence, pilus localization in Gram-positive cocci is determined in both a spatial and temporal manner.

Here we show that surface-exposed *E. faecalis* Ebp are distributed in an asymmetrical manner such that the older hemisphere is saturated with Ebp while new Ebp becomes surface-exposed at the newer hemisphere. This heterogeneous localization pattern has been reported for other *E. faecalis* surface-exposed virulence factors such as aggregation substance (AS) (Olmsted et al., 1993; Wanner et al., 1989). Although localization of surface proteins has been explored in *E. faecalis*, the spatiotemporal distribution of these proteins has not been characterized. Using time-lapse microscopy, we traced the movement of labelled pili along the cell hemisphere and showed that they remain at a fixed position on the cell wall. While positionally fixed on the cell wall, labelled pili are subsequently pushed apart by new cell wall material synthesized at the septum. Ebp turnover appears to be low as labelled pili remain intact throughout cell division. We addressed Ebp turnover by performing a double Ebp immunofluorescence chase experiment using different fluorescently labelled antibodies and observed that new pili are deposited and/or become surface-exposed temporally towards the poles and eventually, the deposited pili converge over the whole cell hemisphere as the pole ages. Hence pilus and AS deposition in *E. faecalis* are coordinated in a specific spatiotemporal manner.

The appearance of septum-exclusive pili on the cell is associated with the age of the cell wall. When different FDAA labels were successively pulsed in short (5 minutes) or long waves (40 minutes) in *E. faecalis*, there was little to no overlap between each label that was pulsed, suggesting there is minimal cell wall turnover, similar to *S. pneumoniae* (Boersma et al., 2015). This finding contrasts with *Bacillus subtilis*, where rapid cell wall turnover is observed when similar FDAA labelling experiments were performed (Boersma et al., 2015; Kuru et al., 2015). The minimal cell wall turnover trait in *E. faecalis* supports the observation that pre-labelled pili remain locked in position as the cell continues to grow and divide. Long pulse chase of FDAA PG labelling in *E. faecalis* showed distinct hemispherical labelling of the older and newer cell wall, indicating that one hemisphere of the cell is always older than the other, as would be expected. This hemispherical labelling pattern is reflective of Ebp localization where surface-exposed Ebp is saturated at one hemisphere of the cell. Together, these similarities led us to speculate that new pili could be intercalated with newly synthesized cell wall. However, our Ebp and cell wall co-staining results showed that previous studies do not fit with the present model. Unexpectedly, the labelling of new PG and surface-exposed pili was almost mutually exclusive, where there was little overlap between the two. When a second Ebp staining was performed after growth in fresh media, new pili appeared to be deposited onto the older pre-labelled cell wall. This result was unexpected as the current paradigm favors that SrtA anchors its substrates onto the cell wall precursor, lipid II, which is subsequently incorporated onto the cell wall via transglycosylation and transpeptidation (Ton-That and Schneewind, 1999; Ton-That et al., 1997). The lack of spatial or temporal correlation between newly synthesised PG and surface-exposed pili suggest that pilus deposition can be independent of lipid II precursors, and that pili can be deposited via an alternative mechanism.

Several lines of evidence involving studies in *S. aureus* indicate that lipid II serves as the cell wall substrate for SrtA. Antibiotic treatment targeting transpeptidation did not affect surface protein anchoring in *S. aureus* (Ton-That and Schneewind, 1999). Furthermore, a mature assembled cell wall was not required for the cleavage of surface protein precursor in staphylococcal protoplasts (Ton-That and Schneewind, 1999). The results of our experiments with *E. faecalis* showed that, with and without inhibition of cell wall synthesis by ramoplanin, an antibiotic targeting lipid II, pilus deposition was not affected. We observed new pili appearing at the cell hemispheres and not at septum where new cell wall synthesis takes place. Based on these observations, it is tempting to speculate that SrtA in *E. faecalis* can anchor cell wall proteins directly onto the crossbridge of uncross-linked, older cell wall. Compared to *S. aureus*, which has approximately 85% of crosslinked PG, the percentage of crosslinked cell wall in *E. faecalis* is lower at approximately 50% (Kim et al., 2014; Yang et al., 2017). The lower percentage of PG crosslinking in *E. faecalis* is attributed to the shorter PG crossbridge of only 2 L-Ala-L-Ala compared to the pentaglycine crossbridge in *S. aureus* (Yang et al., 2017). The abundance of uncross-linked PG in *E. faecalis* may serve as anchor points for SrtA to anchor polymerized pili.

But there’s a conundrum: we see SrtA both at the septum and at the periphery, but only see surface-exposed Ebp at the periphery. We propose at least two possible explanations for this observation. 1) Ebp actually *are* attached to the cell wall at the septum, but they remain inaccessible to antibodies until a later point in the cell cycle or in cell wall maturity. 2) They really are only anchored to hemispherical cell wall, and septal SrtA are inactive for Ebp anchoring. These possibilities are not mutually exclusive. In support of the first explanation, both lysozyme treated WT and sortase mutant cells, in which the cell wall is removed, we observe Ebp puncta on the peripheral membrane of both hemispheres, as well as at the septum. Why then do we not see surface exposed Ebp at the septum? This could be because the cell wall is denser at the septum and more porous at the periphery in an age dependent manner due to the action of the cell wall hydrolases (Wheeler et al., 2015), so Ebp only becomes surface exposed on older cell wall. In support of the second explanation, we observe saturation of Ebp at the cell hemispheres of sortase mutant protoplasts, even when they are only membrane bound. Although we observe Ebp puncta at the septal membrane, it is possible that Ebp are polymerized at the septum, possibly within punctate microdomains, and subsequently move within the cell membrane towards the cell hemisphere for anchoring by SrtA. Microdomains in the bacterial cell membrane are dynamic and fluidic in nature (Los and Murata, 2004; Miller et al., 2019). Furthermore, new surface-exposed Ebp were only seen at the cell hemipsheres with or without cell wall inhibition. The accumulation of membrane bound Ebp at the hemispheres of *ΔsrtA* suggests that the sites of protein cell wall anchoring may also be at hemipsheres. What is not known is whether there is any regulation or coordination involved in guiding membrane bound pili to the hemispheres.

Future work to fully understand how *E. faecalis* Ebp assembly, cell wall anchoring, and surface-exposure is coordinated should include the contribution of cell wall hydrolases. Ebp surface exposure in an *atlA* mutant strain deleted for the major *E. faecalis* cell wall hydrolase (Eckert et al., 2006), occurs at the hemispheres, similar to WT (data not shown). There are at least eight cell wall modifying enzymes encoded in the *E. faecalis* genome (Arthur et al., 1994; Benachour et al., 2012; de Roca et al., 2010; Emirian et al., 2009; Kurushima et al., 2015; Mesnage et al., 2008), and so it would be of interest to determine which (or which combination) might contribute to cell wall porosity, as shown in *B. subtilis* and *S.* aureus (Pasquina-Lemonche et al., 2020) or turnover in *E. faecalis*, leading to Ebp appearance on the old wall. In addition, or alternatively, other cell modifications may be enriched in the more mature PG, such as teichoic acid incorporation or modification (e.g. by alanylation) may favour SrtA substrate incorporation, as has been recently demonstrated in *S. aureus* (Zhang et al., 2021). It is also possible that even though SrtA is localized at the septum, they may be less active for transpeptidation than peripheral SrtA. In *S. aureus*, it has been proposed that the dimeric form of SrtA is more active than the monomeric enzyme (Lu et al., 2007), so perhaps *E. faecalis* SrtA dimerization is favoured within the peripheral membrane.

Taken together, we propose a model where the older hemisphere of *E. faecalis* will always be more saturated with polymerized pili than the younger hemisphere, where newer pili are just beginning to be anchored and exposed to the surface. As the cell grows and elongates during division, new pili are anchored onto uncrosslinked cell wall at the cell hemispheres in sequential order towards the pole and eventually saturate the whole hemisphere (**Fig. 5**). This study provides an in-depth characterization of the spatiotemporal dynamics of peptidoglycan and surface-exposed virulence factors in *E. faecalis* and highlights the possibility of an alternative route to cell wall protein anchoring by SrtA in which substrates are preferentially attached to mature uncrosslinked cell wall rather than lipid II cell wall precursors.

## Acknowledgement

We thank Chris Sham, Kelvin Chong Kian Long, Irina Afonina, Qiao Yuan and Thomas Dean Watts for critical reading of the manuscript. This work was supported by National Research Foundation and Ministry of Education Singapore under its Research Centre of Excellence Programme, as well as the National Research Foundation under its Singapore NRF Fellowship programme (NRF-NRFF2011–11). This work was also supported by a Tier 1 grant sponsored by the Singapore Ministry of Education (MOE2017-T1–001–269). Work in the VanNieuwenhze laboratory was supported by the National Institutes of Health (R35 GM136365). All images were acquired using microscopes from the SCELSE advanced biofilm imaging facility (ABIF). Models in this article were created with the help of BioRender.com.

## Supplementary material

### Aggregation substance chase labelling

*E. faecalis* OG1RF pCF10 cells were grown overnight in BHI supplemented with 15 μg/mL tetracycline and diluted 1:10 in 5mL BHI in the presence of fresh media in the presence of 0.12ng/mL cCF10 peptide to induce aggregation substance (AS) expression for 30 minutes at 37°C, shaking at 200 rpm. Cells were washed once and the first immunofluorescence staining was performed as described previously using rabbit anti-AS serum primary antibody and Alexa fluor 488-goat anti-rabbit secondary antibody (Thermo Fisher Scientific, USA). Next, cells were washed and allowed to grow in fresh BHI supplemented with 0.12ng/mL cCF10 peptide for the second round of AS induction. A second AS immunofluorescence labelling was performed as described but this time, with Alexa fluor 568-goat anti-rabbit secondary antibody (Thermo Fisher Scientific, USA). Cells were then mounted onto glass slides and imaged by SIM.

### Western blot

Cells that were treated and untreated with 26 μg/mL ramoplanin for 1 hour were normalized to OD_600_ 0.5, washed once in PBS and incubated with 10 mg/mL lysozyme in a 37°C water bath for 1 hr. Cell wall and protoplast fractions were separated by centrifugation at 14 000 rpm for 5 minutes. NuPAGE™ LDS sample buffer (Thermo Fisher Scientific, USA) were added to the fractions before boiing for 20 minutes. Ebp blots were run in NuPAGE 3 to 8% Tris-acetate pre-cast gels in Tris-acetate buffer while SecA blots were run in NuPAGE 10% Bis-Tris pre-cast gels for 50 minutes at 120 V. After SBS-PAGE was completed, proteins were transferred onto a PVDF membrane using an iblot™ gel transfer machine and iblot™ transfer stacks (Thermo Fisher Scientific, USA). The membranes were then blocked in 0.1% v/v Tween-5% bovine serum albumin-PBS (Sigma Aldrich, Singapore) overnight at 4°C shaking. The next day, blocking buffer was removed and replaced with primary antibody (rabbit anti-EbpA, rabbit anti-EbpB, guinea pig anti-EbpC and rabbit anti-SecA) at a dilution of 1:3000 for 1 hour at RT and washed trice in 0.01% Tween-PBS. After washing, the membranes were incubated in secondary antibody (IgG guinea pig or rabbit conjugated horseradish peroxidase) (Thermofisher Scientific, Singapore) at a dilution of 1:6000 for 1 hr and washed trice in 0.01% Tween-PBS. Protein bands were detected by chemiluminecence using SuperSigmal™ west femto maximum sensitivity substrate kit (Thermofisher Scientific, USA).

---

**Table S1:** Table of strains and plasmids

| Strain | Resistance<sup>a</sup> | Description | Reference or source |
| --- | --- | --- | --- |
| OG1RF | Rif, Fus | Ebp+, SrtC+, SrtA+, AS– | (Dunny et al., 1978) |
| OG1RFΔ*srtA* | Rif, Fus | Ebp+, SrtC+, SrtA–, AS– | (Guiton et al., 2009) |
| OG1RFΔ*srtC* | Rif, Fus |  | (Nielsen et al., 2012) |
| OG1RFΔ*ebpABC* | Rif, Fus | Ebp–, SrtC+, SrtA+, AS– | (Nielsen et al., 2012) |
| OG1RF/pCF10 | Rif, Fus, Tet | Ebp+, SrtC+, SrtA+ , AS+ | (Afonina et al., 2018) |
| **Plasmid** |  |  |  |
| pGCP123 | Kan | Parent plasmid | (Nielsen et al., 2012) |
| pGCP123 PsrtA SrtA-2L-mCherry | Kan | mCherry tagged SrtA with 2 amino acid linker | This study |

<sup>a</sup>Rif, rifampin; Fus, fusidic acid; Tet, tetracycline; Kan, kanamycin

**Fig S1.**
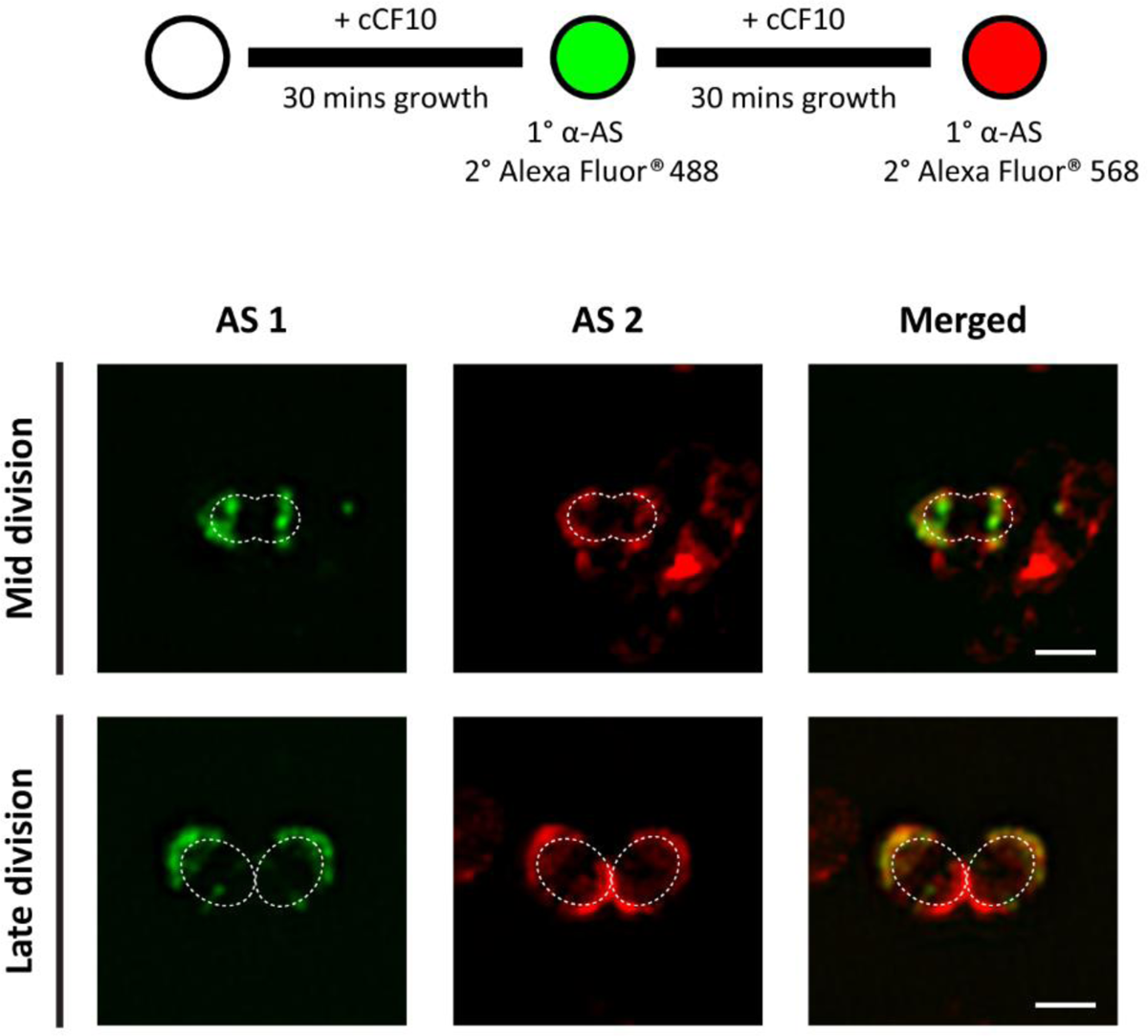
Aggregation substance has similar distribution pattern as Ebp. Chase labelling of AS via immunofluorescence at 30 minutes post induction with cCF10 peptide using green and red fluorescent conjugated secondary antibodies as shown in the schematic where. Representative mid and late division phase cells are shown. Scale bar: 1 µm

**Fig S2.**
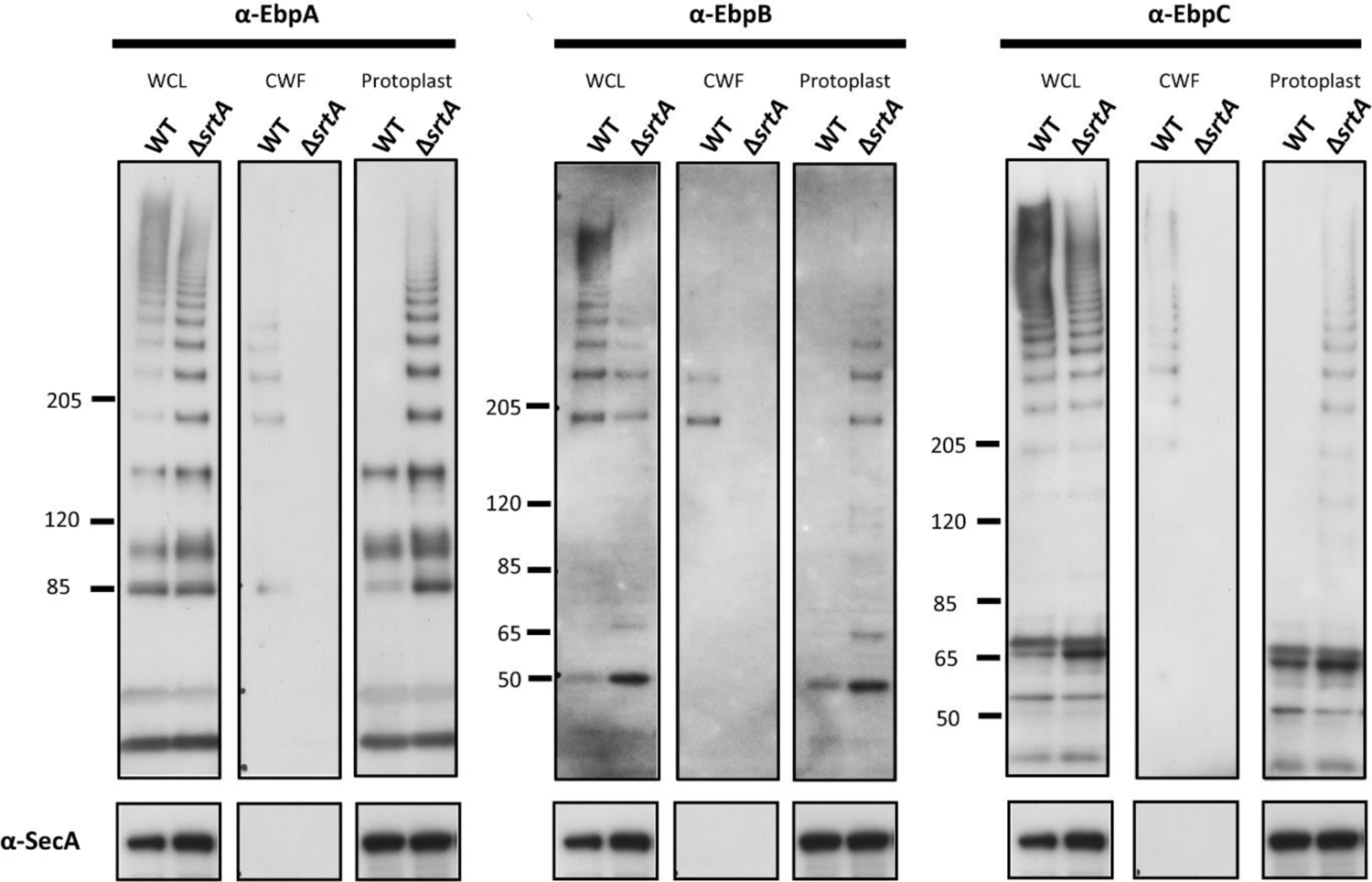
Pili on WT cells are cell wall anchored while pili on Δ*srtA* are membrane anchored. Ebp immunoblots of WT and Δ*srtA* cells in three fractions – whole cell lysate (WCL), cell wall fraction (CWF) and protoplast fraction. All three Ebp subunits were blotted with SecA as a loading control. The predicted size of EbpA, EbpB, and EbpC monomers are 128 kDa, 53 kDa and 68 kDa respectively.

**Fig S3.**
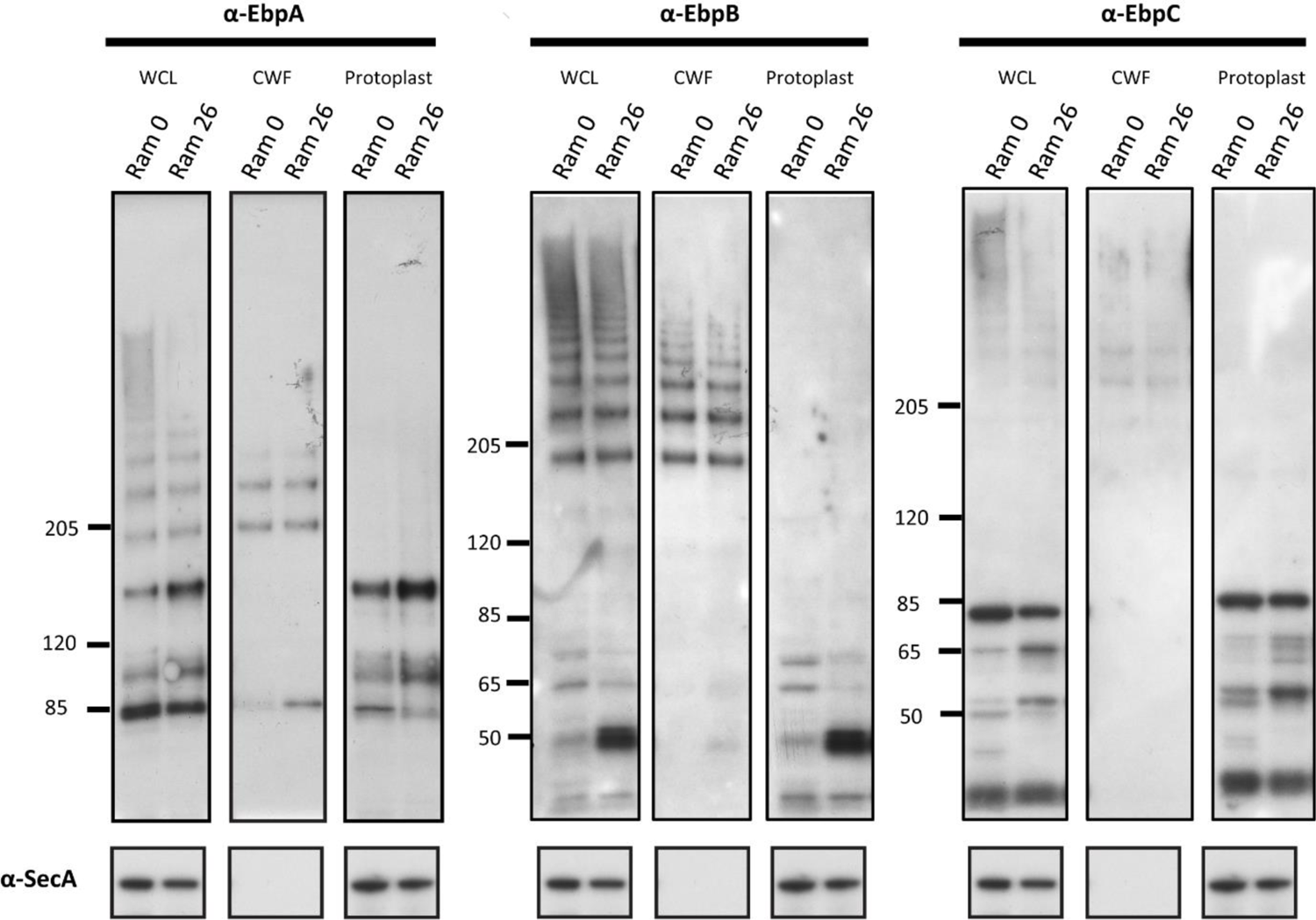
New pili on ramoplanin treated cells are cell wall bound. Ebp immunoblots of ramoplanin treated and untreated cells in three fractions – whole cell lysate (WCL), cell wall fraction (CWF) and protoplast fraction. All three Ebp subunits were blotted with SecA as a loading control. The predicted size of EbpA, EbpB, and EbpC monomers are 128 kDa, 53 kDa and 68 kDa respectively.

